# Comparative population genomics reveals convergent adaptation across independent origins of avian obligate brood parasitism

**DOI:** 10.1101/2025.04.08.647876

**Authors:** Ekaterina Osipova, Christopher N. Balakrishnan, Claire N. Spottiswoode, Jess Lund, Jeffrey M. DaCosta, Mark E. Hauber, Wesley C. Warren, Michael D. Sorenson, Timothy B. Sackton

## Abstract

Parental care evolved as a strategy to enhance offspring survival at the cost of reduced fecundity. While most birds provide parental care, obligate brood parasites circumvent this tradeoff by exploiting the parental behavior of other species. This radical life history shift has occurred independently seven times in birds, offering an outstanding opportunity to test for convergent patterns of adaptation in the genome. To investigate genomic adaptations underlying this transition, we analyzed population resequencing data from five brood-parasitic species across three independent origins of brood parasitism — three parasitic finches (family Viduidae), a honeyguide, and a cowbird — alongside related non-parasitic (parental) outgroups. Using the McDonald-Kreitman framework, we found evidence for repeated adaptation in genes involved in spermatogenesis and sperm function in multiple parasitic clades, but not in the matched outgroup parental species. This is consistent with evidence for increased male-male competition in parasitic lineages as a result of the loss of parental care. In addition, we detected selective sweeps near genes associated with nervous system development in parasitic lineages, perhaps associated with improved spatial cognition that aids brood parasites in locating, monitoring, and laying eggs in host nests. Finally, we found more selective sweeps in host-specific brood parasites as compared to parental outgroups, perhaps reflecting ongoing host-parasite coevolutionary arms races.

## Introduction

The vast majority of birds provide parental care, a reproductive strategy that involves behavioral investment of parental energy and resources to increase offspring survival^1^. However, about 1% of bird species are obligate brood parasites and have foregone parental care altogether, instead laying their eggs in the nests of other species^2^. This strategy means that brood parasites avoid the energetic costs of parental care, from nest building and incubation to feeding nestlings and fledglings. Obligate brood parasitism has evolved seven times in birds: three times in the cuckoos (Cuculidae), and once each in parasitic finches (Viduidae), blackbirds (Icteridae), honeyguides (Indicatoridae), and ducks (Anatidae)^3,4^. These independent origins allow us to investigate the genomic signatures of adaptation associated with a dramatic shift in life history strategy, and to test for genetic evidence of convergent evolution of traits important to a brood-parasitic life history.

Several foundational adaptations associated with the transition from high levels of parental investment to a parasitic lifestyle are likely to be shared by most or all brood-parasitic lineages. As one of the fundamental advantages of brood parasitism, selection for increased fecundity in females is a likely universal adaptation in obligate brood parasites^3,5,6^. As females are freed from the costs of incubation and parental care, time and energetic resources can be reallocated to egg production, and indeed some parasitic species are capable of laying up to ten times as many eggs per season as related parental (non-parasitic) species^7^. Likewise, brood parasites may experience intensified competition between males for mating partners, particularly as compared to species with biparental care, because they are no longer constrained by the time and energy costs of raising offspring. This may lead to increased sexual selection on brood-parasitic males^8,9^. In several *Vidua* finches, for example, males have evolved highly elaborate breeding plumage and/or complex song repertoires^10^, and their mating system has been described as an exploded lek with highly skewed male mating success^11^. In *Molothrus* cowbirds, although female-male consortship occurs, more elaborate plumage and singing behavior in males suggest the influence of sexual selection^12,13^.

Brood parasites do not need to build nests and incubate eggs, but they do need to locate suitable host nests. These nests must not only belong to an appropriate host species (or the “correct” host for a host-specialist parasite), they also must be at the correct incubation stage; parasitising a nest before the host starts laying or after incubation begins often results in parasitism being unsuccessful, though some parasites have evolved tactics to prolong host incubation^2,14^. Therefore, brood parasites have likely experienced selection for enhanced spatial cognition abilities that enable them to locate, remember, and monitor potential host nests^15^.

While brood parasitism allows many of the costs of parenthood to be circumvented, it also introduces a new risk: many hosts have evolved to recognize and reject parasitic eggs and/or chicks^16,17^. Their brood parasites have, in turn, evolved strategies to avoid this host resistance, leading to coevolutionary arms races between increasingly mimetic brood parasitic propagules (i.e., eggs and/or chicks) and increasingly discriminating hosts^18,19^. For example, the cuckoo finch (*Anomalospiza imberbis*) appears to be in a dynamic and ongoing egg-mimicry/rejection arms race with one of its common hosts, the tawny-flanked prinia (*Prinia sublava*)^19,20^. Indigobird and whydah (*Vidua* spp.) nestlings mimic the mouth markings of their respective host chicks^21,22^, and some Australian cuckoos mimic the overall appearance and vocalizations of host young^17,23,24^. The specific adaptations associated with coevolutionary arms races between parasites and hosts vary among parasitic lineages and thus are likely to involve different sets of genes in each parasitic lineage.

Despite the recurrent evolution of this fascinating life history shift, little is known about the genomic underpinnings of the transition to brood parasitism or subsequent adaptations to a parasitic lifestyle. Earlier genomic studies primarily focused on single parasitic lineages or candidate genes related to parental care in mammals. For example, comparative transcriptomic approaches revealed broad-scale differences in neural gene expression patterns between parasitic cowbirds and parental blackbirds^25^, as well as evidence in cowbirds for an insensitivity to prolactin (a hormone often linked to parental behavior)^26^. However, a broad comparative framework is required to test whether the convergent loss of parental care and/or the evolution of derived traits shared across brood-parasitic lineages traces to convergent patterns of change in the genome.

In this study, we conducted a comparative population genomic analysis across three independent origins of brood parasitism, examining both parasitic species and closely related parental outgroups. Using population resequencing data, we identified primary targets of selection across the genome, providing insights into the genomic architecture underlying this evolutionary transition.

## Results

### 1. Resequencing of brood parasites and parental outgroups

To investigate the genomic changes associated with the evolution of obligate brood parasitism in birds, we analyzed population resequencing data for five brood-parasitic species together with related, parental (non-parasitic) outgroup species (table S1). Our sample set included five brood parasites representing three independent origins of brood parasitism, including 16 pin-tailed whydahs (*Vidua macroura*; hereafter ‘whydah’), 19 Cameroon indigobirds (*Vidua camerunensis*; hereafter ‘indigobird’), and 20 cuckoo finches (*Anomalospiza imberbis*) — all members of the parasitic finch family *Viduidae* — along with 31 greater honeyguides (*Indicator indicator*; hereafter ‘honeyguide’), and 10 brown-headed cowbirds (*Molothrus ater*; hereafter ‘cowbird’) (Fig. 1A). Brood parasitism emerged around 22 million years ago in honeyguides, 15 million years ago in finches, and just three million years ago in cowbirds (Fig. 1A) (Carroll et al., in prep). This variation in the age of origin allows us to examine the evolutionary trajectory of brood parasitism by comparing more recent and relatively ancient origins of the trait^27^. To compare genomic signatures of selection between parasitic and non-parasitic groups, we analyzed resequencing data for a species representing a parental outgroup for each of the parasitic groups: the long-tailed finch (*Poephila acuticauda*, n=16; outgroup for parasitic finches), the downy woodpecker (*Dryobates pubescens*; hereafter ‘woodpecker’; n=10; outgroup for honeyguide), and the red-winged blackbird (*Agelaius phoeniceus*; hereafter ‘blackbird’; n=10; parental outgroup for the cowbird)^28^.

**Figure 1.**
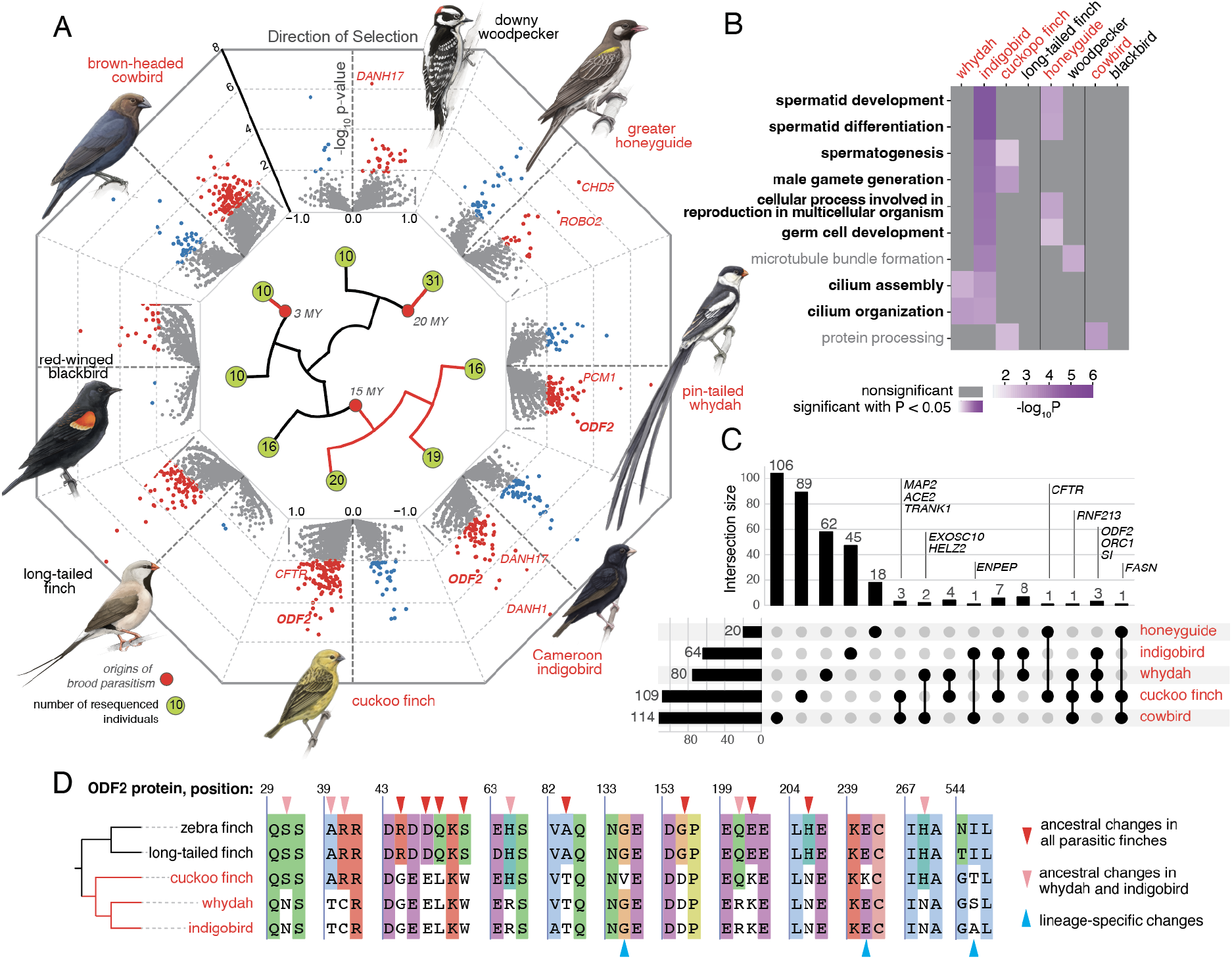
McDonald-Kreitman test revealed positive selection in sperm-associated genes in brood parasites. (A) Tree showing brood parasites and parental outgroups included in the study. Volcano plots visualize the direction of selection (DoS) in the McDonald-Kreitman analysis plotted against -log10 p-value. Genes above the significance threshold (adjusted p-value < 0.01) are highlighted in red (positive DoS) and blue (negative DoS). Red circles at the tree nodes highlight the approximate time of origin of brood parasitism in each clade. The number of resequenced individuals per species is indicated in green circles. Bird drawings by Javier Lazaro. (B) Gene Ontology Biological Processes that are significantly enriched (p-value < 0.05) in the set of positively selected genes in at least two tested species. Purple color indicates significance. Terms relevant to sperm development are in bold; terms significant in at least one of the parental outgroups are in gray. (C) Upset plot illustrating the total number of genes under positive selection in the five brood-parasitic species (horizontal bars on the left) along with the numbers of unique and shared genes (vertical bars). (D) Selected positions in the ODF2 protein, illustrating amino acid substitutions in the ancestor of all parasitic finches (red arrows), the ancestor of the two *Vidua* species (pink arrows), and lineage-specific substitutions (blue arrows).

We called single-nucleotide polymorphisms (SNPs) with the snpArcher pipeline^29^, which maps resequencing reads to a reference genome and calls variants with GATK^30^. The total number of SNPs identified ranged from 11.2 million in indigobird to 20.6 million in cowbird among the brood parasites, and between 22.7 million in woodpecker and 38.4 million in blackbird among the parental outgroups (table S1). This pattern was also reflected in estimates of genetic diversity, which tended to be lower in brood parasites (nucleotide diversity of 0.3% in whydah, indigobird, and honeyguide; 0.4% in cuckoo finch; 0.7% in cowbird) compared to parental outgroups (0.7% in woodpecker; 0.9% in long-tailed finch; 1.2% in blackbird) (table S1). The average read depth across SNPs ranged from 12.8 in whydah to 26.6 in indigobird, indicating sufficient coverage to reliably call SNPs (fig. S2,3, table S1).

### 2. Positive selection in protein-coding genes has targeted reproductive function in brood parasites

To explore patterns of adaptive evolution in protein-coding genes, we used the McDonald-Kreitman test (MKT). This framework contrasts levels of genetic polymorphism and divergence at neutral and functional sites, allowing an inference of positive selection when a larger proportion of nonsynonymous (amino acid changing) fixed differences are observed^31^. The standard implementation of MKT can be biased when the assumption that all polymorphisms observed in a sample are effectively neutral is not met. Given the substantial evidence that weakly deleterious nonsynonymous polymorphisms segregate at appreciable frequencies, several extensions to the standard MKT have been proposed to relax the assumption of strict neutrality^32–34^. We used an imputed MKT extension, which estimates the proportion of weakly deleterious polymorphisms among the low frequency variants and excludes them so that they do not affect estimates of neutral polymorphism^35^. Finally, to quantify and visualize the relative excess of nonsynonymous fixed differences across the five brood parasites, we applied a derivative metric of the direct counts of polymorphisms and fixed differences, the direction of selection (DoS)^36^, where DoS > 0 detects genes with an excess of nonsynonymous fixed differences and signifies positive selection, and DoS < 0 indicates excess segregation of nonsynonymous polymorphisms, which may reflect either relaxation of selection or balancing selection.

Our genome-wide screen for positive selection using the imputed MKT identified 20, 64, 80, 109, and 114 genes under positive selection (FDR adjusted p-value < 0.01 and DoS > 0) in the honeyguide, indigobird, whydah, cuckoo finch and cowbird, respectively. In the three parental outgroups, we detected 28, 95 and 34 genes under positive selection in woodpecker, long-tailed finch, and blackbird, respectively (table S3). We found fewer genes with DoS < 0 (FDR adjusted p-value < 0.01), with brood parasites having between 28 genes (whydah) and 51 genes (indigobird), and parental outgroups having only two genes (long-tailed finch and blackbird) or seven genes (woodpecker) with DoS < 0 (table S3).

Fixed amino acid differences producing the signals of positive selection identified by the MKT could have been inherited from a common brood-parasitic ancestor in each lineage, or arisen independently in each species. In order to determine which type of signal it identified, we compared results for the most closely related brood parasites, whydah and indigobird. In our analysis, we count fixations along the entire branch from the last common ancestor shared with the parental outgroup (the branches shown in red in Fig. 1A). Since whydah and indigobird share a substantial ancestral branch, we expect and also observe counts of fixed differences (D(n) and D(s)) to be correlated between these species (fig. S6). However, we do not find a significant correlation in the direction of selection (DoS) or the MKT p-value (fig. S6). This strongly implies that it is lineage-specific fixations on terminal branches, not shared ancestral fixations, that are responsible for the observed MKT signals in these species, and explains why we see only a small number of shared genes under positive selection. Only three genes show significant evidence of positive selection in all three parasitic finches, whereas an additional eight, seven, and four genes yield significant results in each of the three different pairs of parasitic finches (Fig. 1C). This is consistent with other studies evaluating the power and utility of the MKT; although it relies on fixed differences and therefore is designed to detect selection over relatively deep evolutionary timescales, it primarily detects lineage-specific signals rather than ancestral signals from deeper in time^37,38^.

Several genes identified by the MKT as being under positive selection are plausible candidates for the life history changes associated with brood-parasitism. *SYCP2L* and *CAPRIN2*, detected under positive selection in whydah, are linked to egg production^39^. *BCO1*, which encodes beta-carotene oxygenase 1, is under positive selection in cuckoo finch and whydah. This gene is linked to yellow and orange coloration^40^, and is perhaps involved in the yellow plumage of the cuckoo finch and/or the red bill of pin-tailed whydahs.

To investigate which biological processes are affected by positive selection in brood parasites, we performed a gene ontology enrichment analysis. We found that pathways enriched for genes under positive selection in brood parasites often included those related to sperm development and function (Fig. 1B). We observed this enrichment in whydah, indigobird, cuckoo finch, and honeyguide. We also found significant results for broadly defined cilium assembly and organization terms. In vertebrates, only a few cell types rely on cilia, including those in the respiratory, sensory (primarily mechanoreception, taste, audition, and vision^41^), and reproductive systems. In the reproductive system, cilia are essential for forming the sperm flagellum^42^. Additionally, specific genes enriched for cilia-associated terms in whydah and indigobird, such as *ODF2, DZIP1* and *DNAH8* in whydah and *ODF2, DNAH1* and *DNAH17* in indigobird (table S5), play direct roles in sperm development. This suggests that the cilia-related terms in brood parasites may also indicate enrichment related to spermatogenesis.

Among the genes under positive selection, *ODF2* – which encodes the outer dense fiber protein 2, an essential component of sperm tail structure that is crucial for sperm motility^43^ – was identified in all three parasitic finches (whydah, indigobird, and cuckoo finch; Fig. 1A,C), suggesting that excess fixations on the ancestral branch shared by all three species (Fig. 1D) drives the signal of adaptive evolution in this gene, distinguishing it from the evidence of convergence we observed at the pathway level across distant brood-parasitic lineages (i.e., parasitic finches and honeyguide). Additionally, *DNAH1* and *DNAH17*, which show signals of positive selection in indigobird (Fig. 1A), encode dynein heavy chain proteins that are critical for sperm flagella movement and ciliary function^44–46^. Four other genes (*SI, FASN, RNF213*, and *ORC1*), each of which show evidence of positive selection in three different parasitic species (Fig. 1C), have varied functions unrelated to sperm development, suggesting other axes of convergent selection in brood parasites.

Selection on genes putatively associated with spermatogenesis was identified in all parasitic finches (whydah, indigobird, and cuckoo finch) and honeyguide, which suggests this is a convergent adaptive response in independently evolved brood-parasitic lineages. These findings are consistent with the hypothesis that reduced parental investment in brood parasites intensifies selection on male mating success, increasing male-male competition and producing adaptive changes in genes related to sperm development.

We did not observe enrichment for sperm-associated genes in cowbird, which may reflect differences in the intensity of male-male competition among brood-parasitic lineages. In contrast to many other brood parasites, cowbirds exhibit a consortship-based mating system wherein males associate closely with females, potentially reducing opportunities for multiple mating and thus sperm competition^47^. Notably, cowbird exhibits the largest total number of genes under selection among all the taxa in our analysis. This suggests that the absence of enrichment related to spermatogenesis is not due simply to reduced selection in general, but rather to other genomic targets being under selection.

To address whether the observed enrichment for sperm-associated genes is specific to brood parasites or indicative of a broader trend of rapid evolution in sperm-related genes, we conducted a symmetrical analysis with the parental outgroup species as the focal taxa: long-tailed finch, blackbird, and woodpecker (fig. S5). Our analysis of these parental outgroups did not reveal any significant association with sperm-related processes (Fig. 1B, fig. S9). The lack of similar trends in non-parasitic birds supports the conclusion that selection on sperm-related genes in brood-parasitic species is shaped by their unique life history shift, rather than reflecting a general and recurring trend of rapid evolution in sperm-related genes across most or all bird species. Our results provide evidence of convergent adaptive evolution in sperm development genes associated with the transition to a brood-parasitic lifestyle, suggesting that this is an important axis of selection in parasitic birds.

### 3. Parasitic finches and honeyguide have a higher number of selective sweeps

To search for evidence of recent selective sweeps across the genomes of brood parasites and outgroups, we used SweepFinder2^48^. SweepFinder2 leverages site frequency spectra within genomic windows to distinguish regions that are likely to have evolved under genetic hitchhiking from those that have evolved neutrally. By applying this method across each chromosome, we generated likelihood ratio profiles across the genome (fig. S12A). Given an expectation of an appreciable number of false positives ^49^, we also identified regions of the genome with significant dips in genetic diversity, as measured by nucleotide diversity (*π*) (see Methods for details). We define “sweeps” as regions of the genome with an observed peak in the SweepFinder2 likelihood ratio overlapping a reduction in *π* relative to the flanking regions (fig. S12A).

We consistently identify more sweeps using this approach in honeyguide, indigobird, whydah, and cuckoo finch than in their parental outgroups (Fig. 2A). For parasitic finches, we identified 154 sweeps in cuckoo finch, 165 in indigobird, and 257 in whydah, compared to just 28 in long-tailed finch. Similarly, we identified 168 sweeps in honeyguide compared to 32 in woodpecker. Cowbird, in contrast, did not appear to have more selective sweeps than their parental outgroup. We identified only 12 sweeps in cowbird, which was similar to the 17 we identified in blackbird.

**Figure 2.**
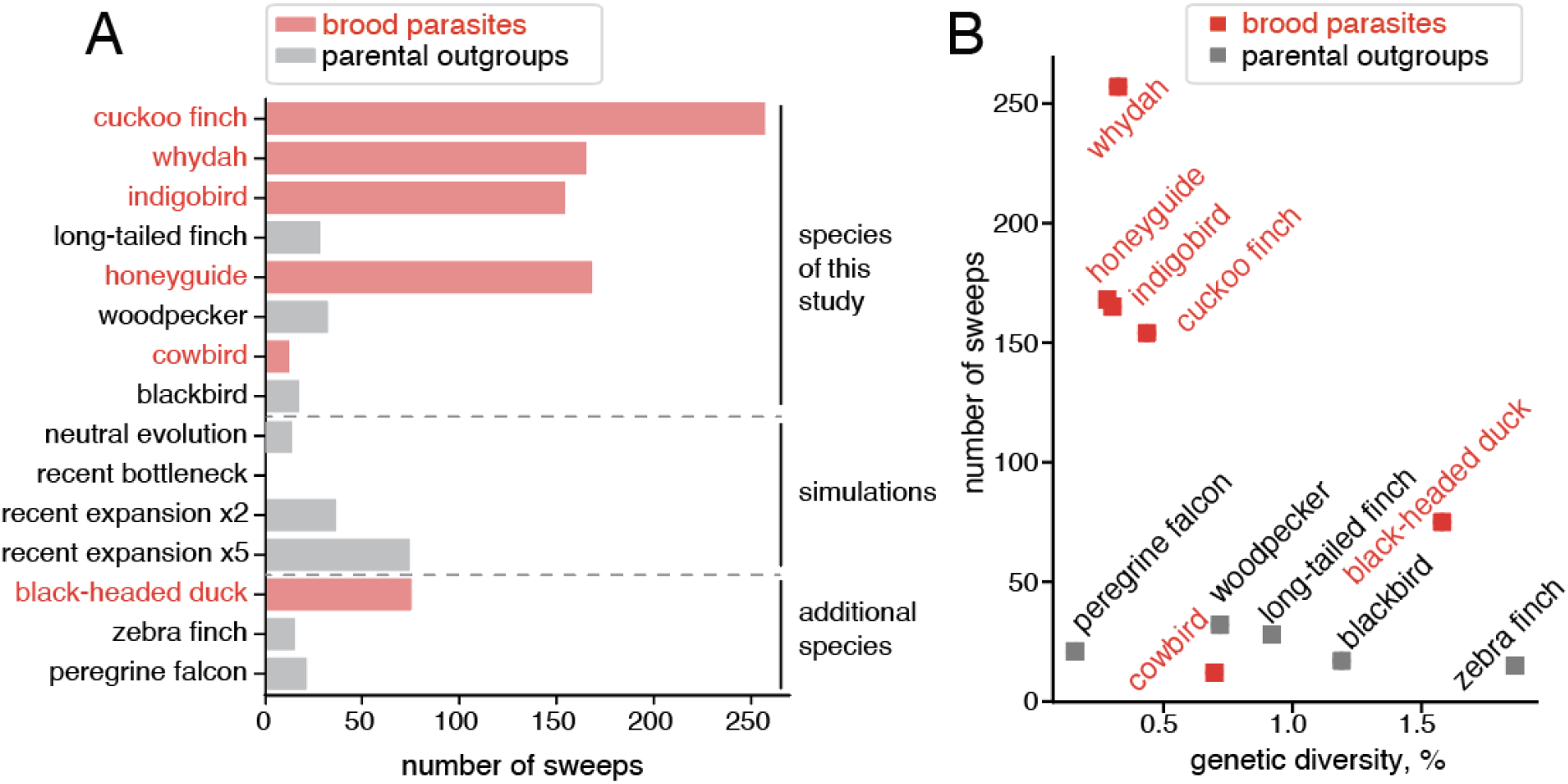
Number of detected selective sweeps is higher in parasitic finches and honeyguide. (A) Total number of selective sweeps detected with SweepFinder2 corroborated by low genetic diversity for brood parasites, parental outgroups, simulated data, and three additional species. (B) Weak negative correlation between the number of selective sweeps detected with SweepFinder2 and genetic diversity computed for the whole genome per species.

To explore the factors contributing to the method’s ability to detect sweeps, we simulated populations evolving under neutrality with different demographic histories (see Methods for details). We found that a recent population bottleneck does not increase the number of detected sweeps, whereas a recent population expansion increases it slightly (Fig. 2A). However, there is no evidence that all parasitic finches and honeyguide have undergone recent population expansions. A recent study suggests that parasitic finches have varied demographic histories, with cuckoo finch and pin-tailed whydah experiencing gradual declines over the past 50–200 ky, whereas Cameroon indigobird shows evidence of recent (<50 ky) population expansion (Sorenson et al., in prep). This suggests that the sweeps we detected and the overall pattern of sweep counts in brood parasites versus parental outgroups was not solely driven by differences in demographic histories.

Since we found that differences in demographic histories could not explain the difference in the number of detected sweeps, we hypothesized that persistent differences in effective population size could explain the observed pattern. Host specialist brood parasites often have smaller population sizes than their hosts^50^, reflecting their dependence on the host species for reproduction^18^. To test this hypothesis, we expanded our selective sweeps analysis to include three additional species: the black-headed duck (a brood parasite with relatively large population size), the zebra finch, and the peregrine falcon. The latter two are parental species with dramatically different population sizes — peregrine falcons have small populations while zebra finches have very large ones. In both peregrine falcon and zebra finch, we detected a number of sweeps comparable to other parental outgroups (Fig. 2A,B). Overall, in tested species, we did not find any significant correlation between genetic diversity – which reflects long-term effective population size – and number of sweeps (Fig. 2B). This suggests that simple population size differences do not explain the observed patterns of sweep detection.

### 4. Recent selective sweeps in brood parasites are near genes involved in neuronal development

Only a small fraction of genomic regions with evidence of a recent selective sweep overlapped coding exons (1% in honeyguide and cuckoo finch, 3 and 4% in indigobird and whydah, respectively), suggesting that selection more often targeted putative regulatory regions. We associated each sweep with the nearest gene (within a 1Mb-domain; see Methods for details). We found that the genes associated with selective sweeps are enriched for neuronal and synaptic development in parasitic finches and honeyguide, but not in cowbird (Fig. 3B, table S8). This suggests that two independent brood-parasitic lineages may be experiencing convergent selection, potentially driven by the demands of their unique ecological niche, which may include enhanced navigation and spatial orientation abilities. To determine if the pattern of selective sweeps near neuronal developmental genes is specific to brood parasites, we again conducted a symmetrical analysis with the three parental outgroup species. None of these outgroups exhibited enrichment in processes associated with synaptic or neuronal function and development, confirming that this pattern is specific to parasitic finches and honeyguide (Fig. 3B).

**Figure 3.**
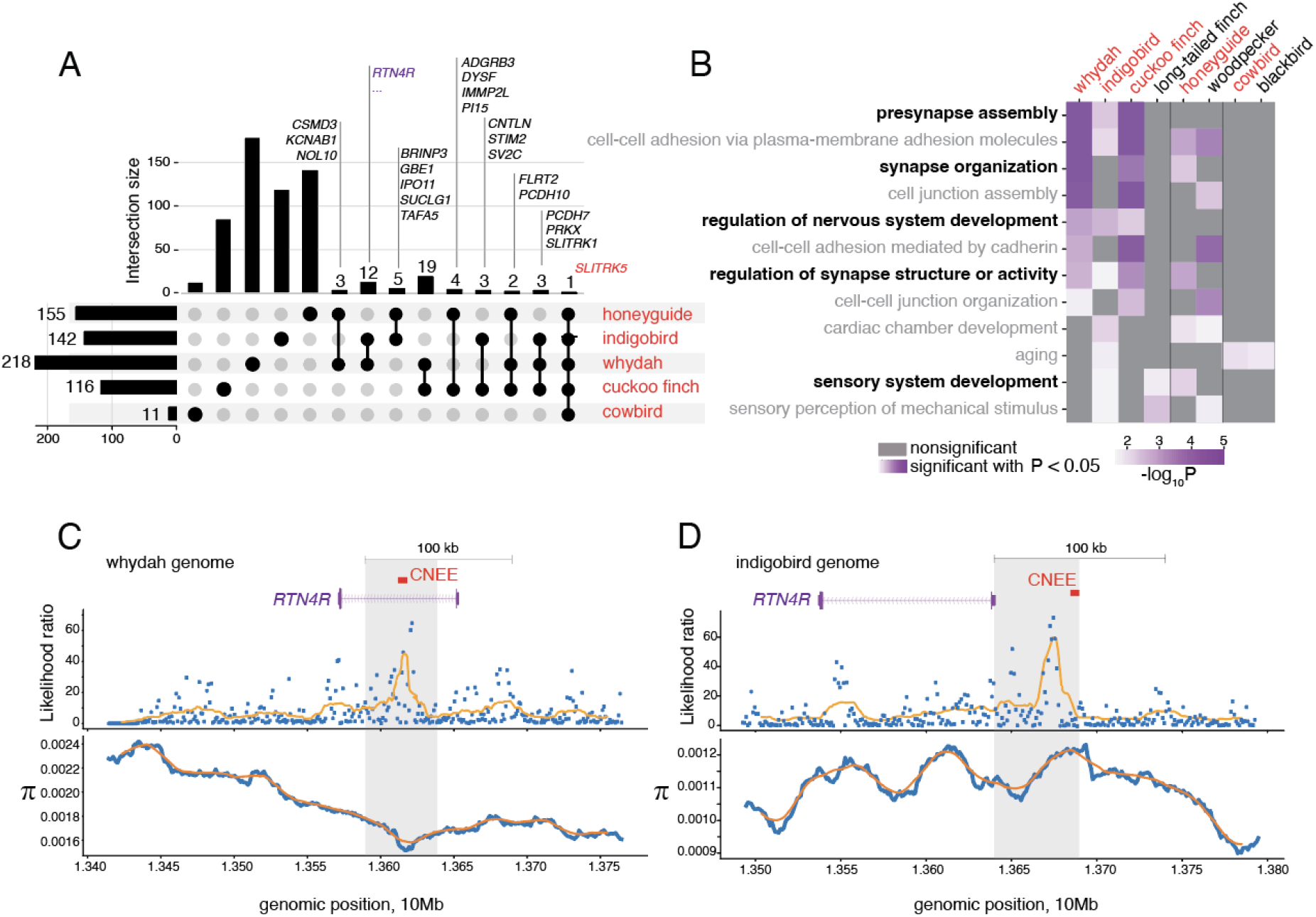
Selective sweeps are located near neuronal development genes in brood parasites. (A) Upset plot illustrating the total number of genes with associated selective sweeps in the five brood-parasitic species (horizontal bars on the left) along with the numbers of unique and shared genes (vertical bars). Genes in sets smaller than ten are indicated, larger gene sets are in table S7. (B) Gene Ontology Biological Processes that are significantly enriched (p-value < 0.05) for positively selected genes in at least two brood parasites. Purple color indicates significance. The terms specific for brood parasites are in bold. The terms that are not specific for brood parasites, but are also present in at least one of the parental outgroups are in grey. (C, D) An example of independent selective sweeps in whydah (C) and indigobird (D) that convergently affect the genomic region encompassing the *RTN4R* gene. The exon-intron structure of the gene is in purple. The overlap between reduced genetic variation (*π*) and likelihood ratio peaks points to selective sweeps in both genomes. Putative regulatory elements (CNEE: conserved non-exonic element) potentially affected by the sweeps are depicted in red.

Some of the genes with associated selective sweeps are plausible candidates for links to the transition to brood parasitism. *NLGN4X* (neuroligin 4 X-linked), which is associated with putative sweeps in both cuckoo finch and whydah, is involved in synaptic function and social behavior and has been linked to parental care^25,51^. Other genes associated with parental care in birds^25^ include *GRPR* (gastrin-releasing peptide receptor), *NTF3* (neurotrophin 3), and *HTR2C*, which encodes a serotonin receptor; these genes are associated with putative sweeps in indigobird, cuckoo finch and honeyguide, respectively. *TGFBR2*, which shows evidence of a selective sweep in indigobird, is a developmental gene also associated with egg production^52^. Finally, *OCA2*, which encodes a melanosomal transmembrane protein^53^, shows evidence of a sweep in both whydah and indigobird. *OCA2* may contribute to the black and/or iridescent plumage colors of viduine finches used for sexual attraction.

At the gene level, we also observed a surprising pattern of convergence in independently evolved parasitic lineages. For example, the *SLITRK5* gene, which is associated with sweeps in four brood parasites (all three parasitic finches and honeyguide, Fig. 3A), encodes a protein with brain-specific expression that plays a crucial role in neurite outgrowth and synapse formation, contributing to the modulation of synaptic transmission^54^. Similarly, *SLITRK1*, associated with sweeps in all parasitic finches, also functions in neuronal development^55^. Protocadherins 10 and 7, encoded by *PCDH10* and *PCDH7*, are associated, respectively, with sweeps in all three parasitic finches, and in cuckoo finch, whydah, and honeyguide. They are involved in cell-cell adhesion processes in the brain, contributing to neural connectivity^56,57^. *FLRT2*, which is associated with sweeps in cuckoo finch, whydah, and honeyguide, also plays a role in cell adhesion and signaling in neural development, contributing to synaptic connectivity^58^. Collectively, the majority of genes associated with sweeps in at least three different brood parasites play a distinctive role in neuronal development and function, reinforcing the patterns observed at the pathway level.

We also detected a compelling example of gene-level convergence between whydah and indigobird. We identified evidence of selective sweeps in both species in proximity to *RTN4R*, which encodes Nogo receptor 1. This is a protein tyrosine phosphatase receptor that has been shown to have an important role in neuronal development and synaptic plasticity, specifically in processes involved in learning and spatial memory^59,60^. Although these species are in the same genus, it is notable that the evidence for selective sweeps in these two species falls within different portions of the genomic region containing the *RTN4R* gene – specifically, in the intronic region of the gene in whydah (Fig. 3C) and upstream of the gene in indigobird (Fig. 3D) – supporting a hypothesis of convergent evolution rather than a shared signal of selection from a common ancestor. In addition, we found that in both cases, the evidence for a selective sweep overlaps conserved non-exonic elements, suggesting that these sweeps target putative regulatory regions that affect expression of the gene. That different portions of the same genomic region show evidence of selection suggests independent bouts of selection in these two species, perhaps related to convergent or divergent selection on the same trait in both species.

## Discussion

We used whole-genome resequencing data from five brood-parasitic bird species and representative parental outgroups to investigate genomic signatures of natural selection associated with brood parasitism. By identifying signatures of selection across three independent origins of this major life history transition, we can discriminate between examples of lineage-specific evolution and convergent changes due to selection on traits generally associated with brood parasitism and the loss of parental care. By applying this comparative approach, we identified convergent patterns of ongoing selection within parasitic lineages, shedding light on the genomic signatures associated with this remarkable behavioral strategy.

Our results suggest that selection on protein-coding genes has primarily targeted reproductive function in brood parasites. Specifically, we found signatures of selection on genes relevant to sperm development and function in honeyguide and parasitic finch, which were not present in their parental outgroups. This likely reflects an increased intensity of male-male competition following the loss of parental care. All the parasitic species we analyzed are sexually dimorphic, with some degree of elaborate male plumages along with courtship vocalizations and visual displays. The two *Vidua* species may be most extreme in this regard, with the indigobird mating system described as a dispersed lek^50^ and pin-tailed whydah having elaborate male plumage and flight displays. Our results are consistent with other studies suggesting that lekking and highly polygynous birds show clear signals of positive selection on genes underlying male display traits, from brilliant plumage coloration to muscle performance^61,62^.

Surprisingly, we did not find similar genomic signatures of selection on sperm-related genes in cowbird. A possible explanation for this is the socially monogamous mating system of brown-headed cowbirds, where males associate closely with females^63,64^. The mate-guarding observed in cowbirds may prevent the female from mating with multiple males, thereby reducing sperm competition after mating and reducing the intensity of selection on sperm-associated genes. Alternatively, the lack of genomic signatures related to sperm development may be due to the relatively recent origin of brood parasitism in this group (approximately 3 million years ago in cowbirds, compared to over 15 and 20 million years ago in brood-parasitic finches and honeyguides respectively, Carroll et al. in prep).

Our results also showed genes related to neuronal development were overrepresented in genomic regions affected by selective sweeps. Some of these genes are directly associated with spatial memory and navigation, which suggests that brain-related traits continue to evolve under positive selection in brood parasites. It is likely that this reflects continuous selection in brood parasites on traits associated with accurately locating and storing information about host nests. In greater honeyguide, selection on cognitive abilities might also be associated with their mutualistic relationship with humans. Uniquely among birds, greater honeyguides guide people to the locations of active bees’ nests, which the people harvest to collect the honey^65^. Honeyguides are then able to access the nutrient-rich wax within the bees’ nest. Storing information about the locations of bees’ nests in an environment is likely a cognitively expensive task, such that greater honeyguides may be under ongoing selection for improved spatial memory in this context. However, the pattern of selective sweeps near neuronal genes is convergent in honeyguide and parasitic finches, which supports the hypothesis that these genetic changes are at least in part associated with the transition to a brood parasitic life history and the shared problem of tracking host nests.

We observed a consistently higher number of selective sweeps in parasitic finches and honeyguide compared to their parental outgroups. One possible explanation for this pattern is that differences in population size or demographic histories of the two groups may influence our ability to detect sweeps. However, our analysis showed no significant correlation between population size and the number of detected sweeps (Fig. 2B). Furthermore, if parasitic finches and honeyguide have more dynamic demographic histories compared to the outgroups and cowbird, for example, due to host switching events, this could influence sweep detection. However, our simulations demonstrate that while population expansions can generate low levels of false positive signal, demographic history alone cannot generate the number of sweeps we see in most parasitic lineages (fig. S11, Fig. 2A). Additionally, the parasitic finches we sampled have different demographic histories over the past million years or more (Sorenson et al., in prep), further weakening this possible explanation. Therefore, the higher number of selective sweeps in two of the brood-parasitic clades suggests that they have experienced more episodes of strong selection than their parental outgroups.

Cowbird showed a drastically lower number of selective sweeps compared to the other four parasitic species we investigated. A potential explanation for this pattern is that cowbirds have a shorter evolutionary history as brood parasites (only 3 million years) (Carroll et al., in prep) and are extreme host generalists, exploiting over 200 host species^66^. Effectively, the lack of long-term coevolutionary interactions with specific hosts may explain the paucity of detectable selective sweeps in cowbird. Parasitic finches and honeyguides, in contrast, adopted brood parasitism over 15 million years ago and show clear evidence of genetic specialisation on different host species or groups of related host species^67,68^. Together our results suggest that host-parasite coevolutionary interactions and the duration of these interactions are key factors in producing the larger number of detected sweeps. However, it is difficult to disentangle whether the pattern of selective sweeps we observed is due specifically to host-parasite coevolution or time since the origin of brood parasitism in each clade.

Overall in this study, we found both convergent and lineage-specific signatures of selection underlying the transition to a brood-parasitic life history. Our findings highlight the importance of evolutionary history, host-parasite coevolutionary interactions, and mating systems in shaping genomic signatures of selection. However, phylogenomic approaches aimed at identifying changes occurring closer to the base of parasitic clades will be required to identify key genes and regulatory changes associated with the loss of parental care behavior.

## Methods

### Reference genomes and population resequencing data

Our analyses are based on several previously available and seven newly assembled whole genome assemblies for both parasitic and parental species. The new genomes (Carroll et al. in prep) include the brood-parasitic cuckoo finch *Anomalospiza imberbis*, village indigobird *Vidua chalybeata*, pin-tailed whydah *Vidua macroura*, brown-headed cowbird *Molothrus ater*, and greater honeyguide *Indicator indicator*, along with the parental species red-winged blackbird *Agelaius phoeniceus* and downy woodpecker *Dryobates pubescens*.

We generated population resequencing data for 31 individuals of the honeyguide species *Indicator indicator* and 10 individuals of downy woodpecker *Dryobates pubescens*. For the other species in our analysis, we used published population resequencing data, which included samples of 10 to 20 individuals (Sorenson et al., in prep). Data for the long-tailed finch, zebra finch, and peregrine falcon were obtained from GenBank, while we produced new data for the remaining species.

### Variant calling

#### Running snpArcher

We used the snpArcher pipeline ^29^ to call variants in population resequencing data for each brood parasite. Using the pipeline, we mapped resequencing reads of each species to the reference genome of that species, with the exception of Cameroon indigobird in which resequencing reads were mapped to the village indigobird genome. All reference genomes were taken from NCBI and the identifiers are listed in table S1. In addition to the GATK best practices filtering ^30^, we filtered the resulting VCF files removing indels, non-biallelic SNPs, SNPs with a minor allele frequency < 0.01, SNPs with >75% missing data, and samples with <2x sequencing depth using bcftools view (v1.10). In addition, we removed reference as one of the ingroups by excluding all SNPs with allele frequency = 1.0 `(−v snps -U -m2 -M2` `-e ‘F_MISSING > 0.75 | AF=1.0 | ref=“N” | ALT=“.” | TYPE∼”indel”‘)`.

#### Quality control and sample exclusion

To evaluate data quality for each sample, we used the snpArcher QC (Quality Control) module on a randomly selected subset of ∼100,000 SNPs. This analysis included principal component analysis (PCA), assessments of missing data, average read depth, and relatedness between resequenced individuals. We excluded individuals with high levels of missing data or those closely related (first- and second-degree relatives). **Pin-tailed whydah**: one sample (P3_JS-39_LH) was excluded due to high missingness and low SNP depth (fig. S2,3). **Cameroon indigobird**: four samples (CNB452, CNB251, CNB250, CNB453) were excluded because – as expected based on sample collection – they clustered separately from the main population on the PCA plot, representing samples of a different species, theWilson’s indigobird (*V*.*wilsoni*) (fig. S1). **Downy woodpecker**: ten samples were excluded based on PCA, which indicated they belonged to a different species, the hairy woodpecker (*L. villosus*) (602078_S39, 643636_S40, 643663_S41, 601316_S42, 627823_S43, 641543_S44, 611737_S45, 633898_S56, 363492_S21, 428786_S22). **Long-tailed finch**: four samples (ERR1013135, ERR1013137, ERR1013145, ERR1013150) were excluded due to being likely second-degree relatives or having >50% missingness (fig. S2,S3).

### McDonald-Kreitman test

To apply the McDonald-Kreitman framework, we constructed trios of species, each trio consisting of ingroup resequenced individuals and two outgroups (fig. S5). Each VCF file of ingroups was polarized with the first outgroup followed by adding the second outgroup. To polarize VCF, we ran whole genome alignment with minimap2 (v2.24-r1122) ^69^ (`minimap2 -cx asm20 --cs`) followed by running paftools (paftools.js call) to convert PAF alignments to VCF. Finally, we filtered the resulting VCF files for SNPs only with bcftools view (v1.10) `(−v snps -U -m2 -M2 -e ‘F_MISSING > 0.75 | ref=“N” | ALT=“.” | TYPE∼”indel”‘)`. To search for genes with an excessive fraction of fixed differences compared to polymorphic changes, we ran Degenotate (https://github.com/harvardinformatics/degenotate). When available, we used RefSeq annotations generated by the NCBI annotation pipeline (table S1). For the long-tailed finch, we ran TOGA ^70^ to project zebra finch *Taeniopygia guttata* RefSeq annotation (GCF_003957565.2) based on the whole-genome alignment of the long-tailed finch genome^28^ to the zebra finch genome (GCF_003957565.2). To mitigate the biases of the standard MKT framework that treats all polymorphisms as effectively neutral, we implemented the imputed MKT extension ^35^ in Degenotate. Degenotate calculates several MKT-related statistics including imputed p-value and imputed direction of selection (DoS) ^36^:

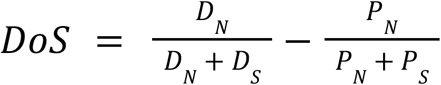

which we used for all downstream analyses. To select top gene candidates, we applied an adjusted p-value < 0.01 cutoff for both positive and negative DoS.

### Screen for selective sweeps

#### Polarization of variants

To prepare input for SweepFinder2, we first polarized VCF files of all target species using genomic data of corresponding outgroups. Each outgroup genome was aligned to the corresponding target genome using minimap2 (v2.24-r1122) ^69^ as described in the McDonald-Kreitman analysis section. The ancestral state was inferred using bcftools consensus (v1.10) to generate a consensus FASTA file from the outgroup VCF. A merged VCF file was created using bcftools merge (v1.10) with missing data set to reference (`--missing-to-ref`) for both the query and outgroup VCF files. Ancestral alleles were inferred with vcfdo polarize (https://github.com/IDEELResearch/vcfdo).

#### Preparation of SweepFinder2 input

VCF data were filtered using vcftools (v0.1.16) `(vcftools --counts2 –derived --vcf`), and each chromosome chunk was processed for SweepFinder2 input by excluding sites with zero frequency. A header was added to ensure proper formatting for SweepFinder2. We divided large chromosomes into 2 Mb segments for further analysis.

#### SweepFinder2 Analysis without recombination map

We ran SweepFinder2 without a recombination map to identify regions under selective sweeps. SweepFinder2 was executed using a log grid size (`-lg 1000`).

#### Recombination map preparation

To account for variation in recombination rates across the genome, we used zebra finch recombination map ^28^ and lifted it to the target genomes using liftOver ^71^ and alignments generated from each target species to the reference zebra finch genome. In the first step, we lifted the zebra finch recombination map to the newer zebra finch genome assembly (GCF_003957565.2). Next, we lifted this resulting recombination map to each of the target brood parasite and outgroup genomes using whole-genome alignments of the target genomes to zebra finch. These maps were further refined using bedtools (v2.30.0) to merge intervals with gaps smaller than 10 bp.

#### SweepFinder2 analysis with recombination map

Using the prepared recombination maps, we reran SweepFinder2 on the same chromosome segments to identify selective sweeps accounting for recombination. The results were combined with those obtained from the non-recombination map analyses.

#### Merging and reconciling results

Results from SweepFinder2 runs with and without recombination maps were combined using bedtools intersect (v2.30.0). This allowed us to integrate the two sets of results and produce a comprehensive output of regions under selection across the genome.

#### Calculating genome-wide Pi and depth

Nucleotide diversity (*π*) was computed genome-wide using vcftools (v0.1.16) (`--window-pi 100000 --window-pi-step 1000`). Genome-wide read depth was calculated for each chromosome, with depth values summed over 1 kb windows. Depth values were computed using samtools depth (v1.17) and a custom script (`compute_window_sum.py`).

#### Integrating SweepFinder, Pi, and depth results

The likelihood ratio (LR) results from SweepFinder2, nucleotide diversity (*π*) calculations, and genome-wide depth were combined using bedtools intersect (v2.30.0). We detected local extrema in *π* and LR applying a simple moving average to smooth the input data, using a sliding window size of 20 kb. The smoothed data were then analyzed to identify local minima (for *π*) and maxima (for LR) using the `find_peaks` function from the python3 *scipy* library. Extrema were detected based on specified prominence thresholds: *π* = 0.0001 and LR = 10. We defined selective sweeps as regions of the likelihood ratio (LR) local maxima overlapping *π* local minima. Identified *π* minima were extended by ±25,000 bp to define candidate selective sweep regions. We excluded all regions with depth below 100 reads yielding a final dataset for downstream analysis of genomic regions with potential selective sweeps.

#### Simulation variation under different evolutionary scenarios

We ran MSPrime^72,73^ under different evolutionary scenarios to simulate VCF files of populations evolving neutrally, under the scenario of recent bottleneck, and under the scenario of recent population expansion using a custom script `simulate_vcf`.`py` (available at https://github.com/osipovarev/brood_parasites). We simulated 500 chromosomes of 2Mb length each (1Gb genome in total, comparable to the average size of a bird genome) for a population of 20 individuals, with an effective population size of 100,000, which is comparable to an average estimated for brood-parasitic population. We used a mutation rate of 1e-8 and a recombination rate of 1e-8. We defined a recent bottleneck as an effective population size change from 100,000 to 1000 individuals, happening between 100,000 and 50,000 generations ago. We defined a recent population expansion as an effective population size change from 100,000 to 200,000 and 500,000 individuals (2x and 5x), happening between 100,000 and 50,000 generations ago. We polarized this VCF with one outgroup that was simulated using the mutation rate 1e-7. Next, we ran SweepFinder2 on the simulated data the same way as we did for the real population resequencing data.

### CNEEs liftOver

#### Modeling and masking repeats

For the genome assemblies of brood parasites and outgroups, we used RepeatModeler ^74^ to identify repeat families and RepeatMasker ^75^ to soft-mask repeats. For other bird genomes, we used the available repeat masker annotations provided by NCBI. All species names and their assemblies are listed in table S1.

#### Generating pairwise genome alignments

To compute pairwise genome alignments between zebra finch *T. guttata* (GCF_003957565.2) as references and other birds as query species, we used LASTZ version 1.04.00 ^76^ with parameters K = 2,400, L = 3,000, Y = 9,400, H = 2,000 and the default scoring matrix, axtChain ^77^, chainCleaner ^78^, and RepeatFiller ^79^ (all with default parameters). For the downstream analyses, we excluded chains with a score of less than 100,000.

#### CNEEs liftOver

To analyze the association between selective sweeps and conserved non-coding elements, we projected CNEEs annotated for *G. gallus* (galGal7) onto *T. guttata*. Next, based on the alignment of each brood parasite and outgroup we projected CNEEs from *T. guttata* coordinates to the coordinates of the query genomes.

### GO enrichment analysis

We performed the enrichment analysis with the R package ClusterProfiler (v4.8.2)^80^ using human OrgDb org.Hs.eg.db and Gene Ontology (GO) Biological Processes subontology. To be able to use the human database, we converted bird gene IDs into human gene IDs through chicken gene IDs as an intermediate step using biomart (https://useast.ensembl.org/biomart/martview). We ran enrichGO analysis applying cutoffs for the size of the searched terms, with minimum size = 40 and maximum size = 400. We excluded the terms with the number of gene hits fewer than 3. All genes annotated for a species were used as a background (gene universe) set. Finally, we selected significantly enriched terms applying p-value cutoff < 0.05.

## Data and code availability

Population resequencing data of the greater honeyguide and downy woodpecker are available on NCBI under the accession (accessions pending).

The code is available at https://github.com/osipovarev/Brood_parasites_analysis

## Competing interests

The authors have no competing interests.

## Acknowledgments

This work was supported by collaborative NSF grants to MDS, CNB, WCW, TBS and JMD (DBI 1754311, 1754397, 1754406, 1754546, 1754643, 1940624) and to MEH and CNB (IOS 1456524, 1456612, 1953226). Funding for fieldwork on honeyguides and cuckoo finches was supported by grants to CNS from European Research Council (Consolidator Grant 725185), Biotechnology and Biological Sciences Research Council (BBSRC, David Phillips Research Fellowship, BB/J014109/1) and the Royal Society (Dorothy Hodgkin Fellowship). For their efforts in the field and laboratory, we thank: Gabriel Jamie, Tanmay Dixit, Silky Hamama, Collins Moya, Wenfei Tong (cuckoo finches); Iahaia Buanachique, Dominic Cram, David Lloyd-Jones, Mussaji Muamedi, Carvalho Issa Nanguar, Jessica van der Wal, Orlando Yassene, Susan Miller (greater honeyguides); Justin Schuetz (pin-tailed whydahs); and Bruce Lyon, John Eadie, Melissa Jones (black-headed ducks). Brown-headed cowbird, red-winged blackbird, and downy woodpecker samples were provided by the United States National Museum of Natural History. We thank Melissah Rowe, Judith Risse, Simon Griffith, and Daniel Hooper for advance access to their *Poephila* genome assembly. We also thank Administração Nacional das Áreas de Conservação, Mozambique, for permission to work in the Niassa Special Reserve, Mozambique. (Permit numbers: 008/2015, 11/11/2016, 15/2019). For assistance and discussions, we thank Matt Louder. Bird drawings were prepared by Javier Lazaro.

## Notes

### Competing Interest Statement

The authors have declared no competing interest.

