## Supplementary Figures for "Comparative population genomics reveals convergent adaptation across independent origins of avian obligate brood parasitism"

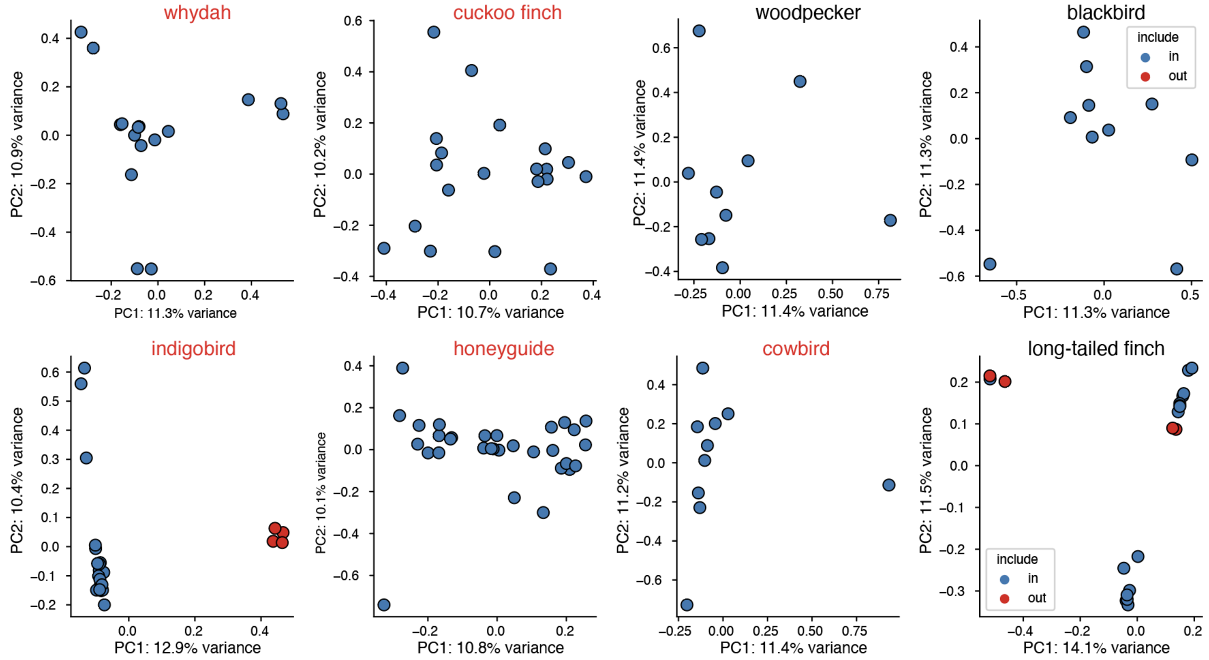


**Fig. S1. PCA plots for each species generated from snpArcher data.**
Blue samples were included from the downstream analyses, red samples were removed. Red dots in the indigobird plot represent samples of another species – Wilson's indigobird (*V.wilsoni*) – which were excluded in the downstream analyses. In the woodpecker data, we additionally excluded samples representing hairy woodpecker (*L.villosus*), not shown here. Red dots in the long-tailed finch plot represent samples excluded due to high missingness (fig. S2) or samples coming from close relatives (fig. S4).

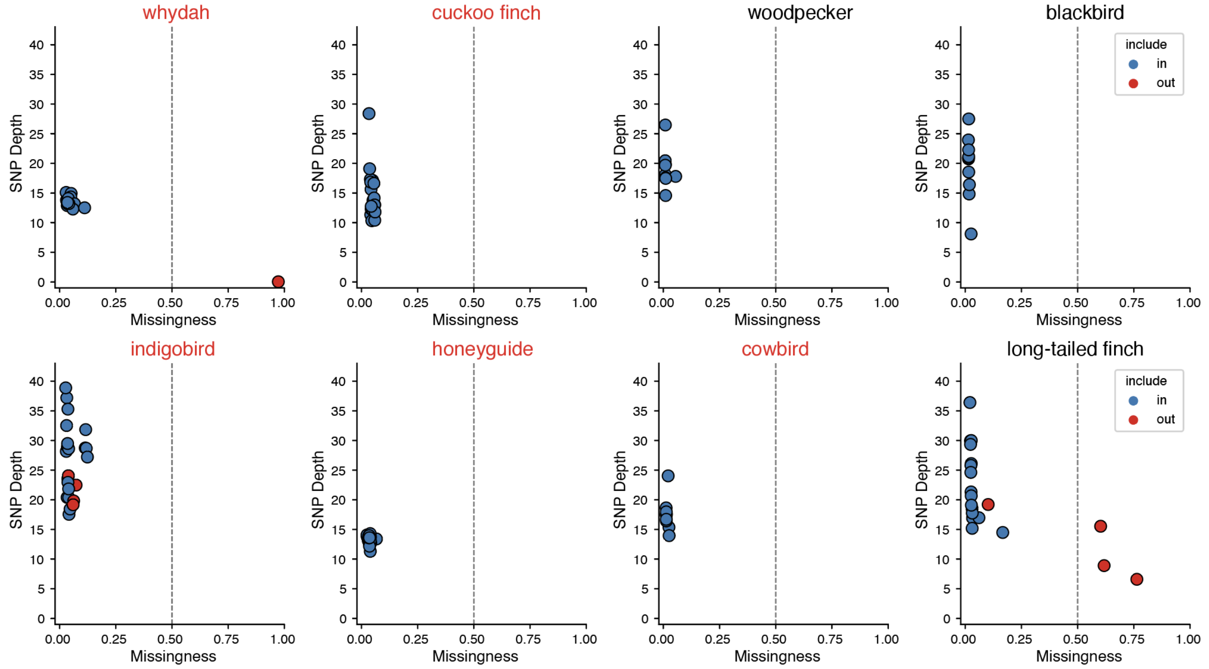


**Fig. S2. SNP depth and missingness for each analyzed species.**
Blue samples were included from the downstream analyses, red samples were removed.

The red dot in the whydah plot represents an excluded sample that had almost no reads mapped to the genome. Red dots in the indigobird plot represent samples of another species – Wilson's indigobird (*V. wilsoni*) – which were excluded in the downstream analyses. Red dots in the long-tailed finch plot represent samples excluded due to high missingness or samples coming from close relatives (fig. S4).

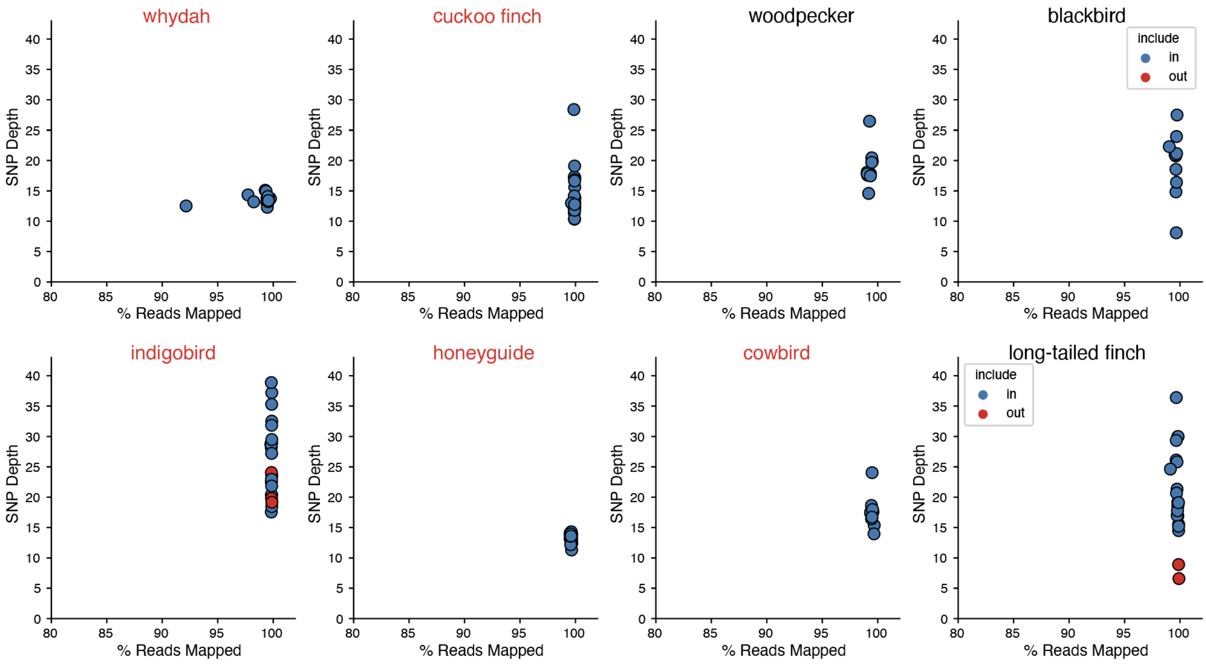


**Fig. S3. SNP depth percentage of reads mapped for each analyzed species.**
Blue samples were included from the downstream analyses, red samples were removed. Red dots in the indigobird plot represent samples of another species – Wilson's indigobird (*V.wilsoni*) – which were excluded in the downstream analyses. Red dots in the long-tailed finch plot represent samples excluded due to high missingness (fig. S2) or samples coming from close relatives (fig. S4).

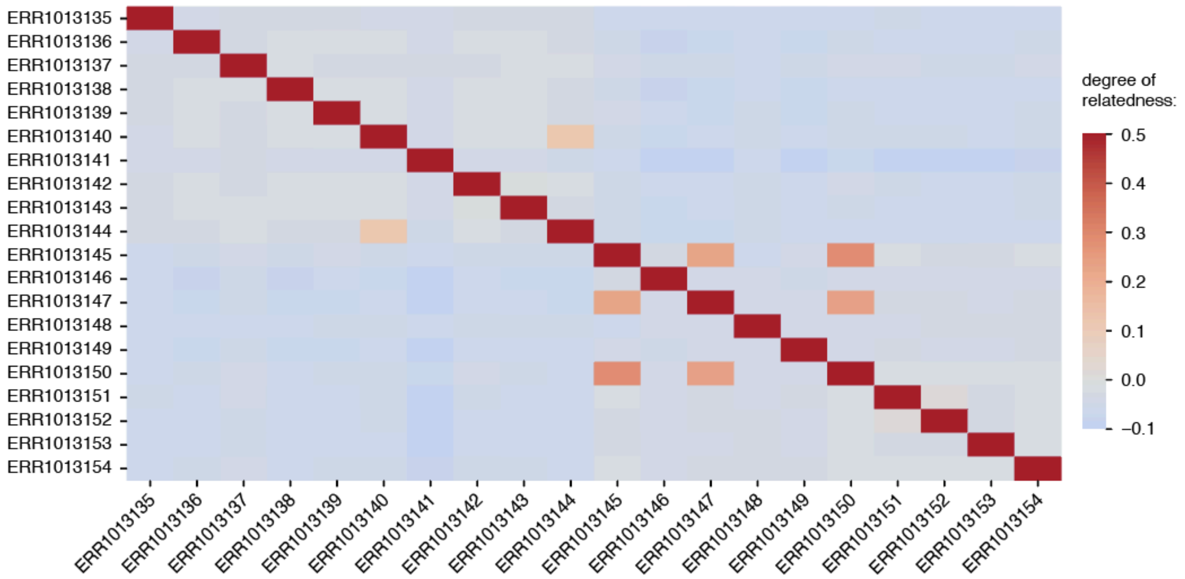


**Fig. S4. Relatedness matrix between resequenced individuals of the long-tailed finch.** Samples with relatedness of 0.5 indicate they are identical, values close to 0.25 indicate parent-child or sibling relatedness and second-degree relatedness is ~0.125. We removed closely related individuals (>0.125) from the downstream analyses.

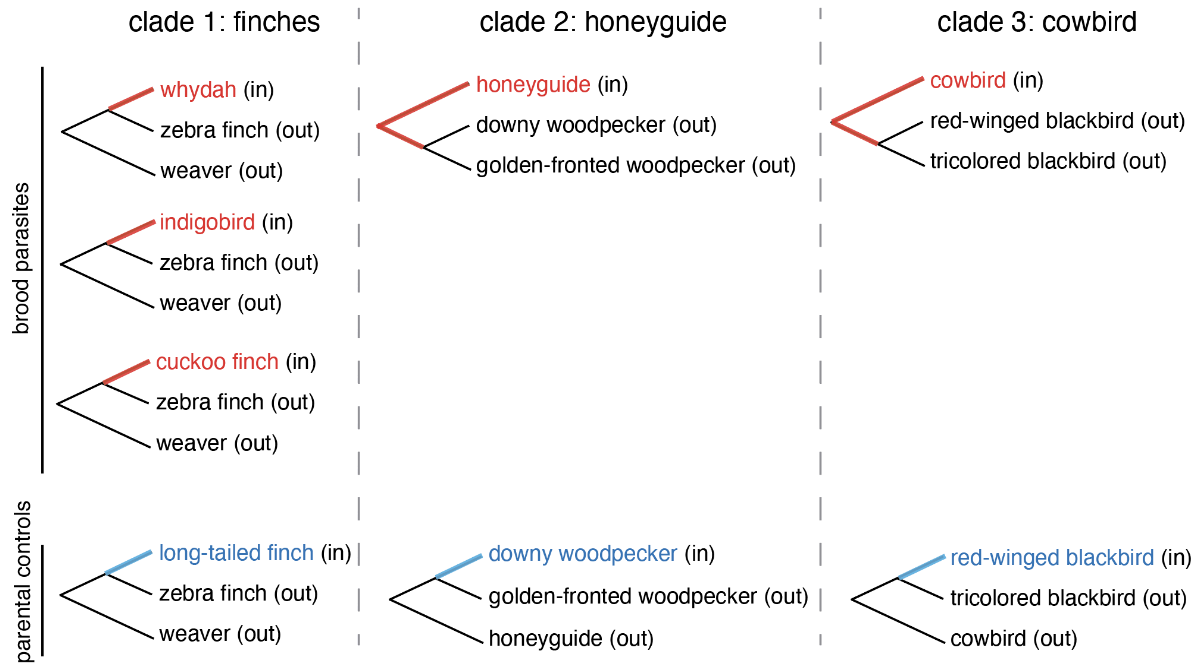


**Fig. S5. Trios of brood parasites and parental outgroups for McDonald-Kreitman test.** (A) Finches clade that includes parasitic finches (whydah, indigobird, and cuckoo finch) and the long-tailed finch outgroup.
(B) Honeyguide clade that includes honeyguide and the downy woodpecker outgroup.
(C) cowbird clade that includes cowbird and the red-winged blackbird outgroup. Among the target species for each analysis, brood parasites are colored in red, and parental outgroups are colored in blue. in=ingroups represented in the analysis with resequenced individuals; out=outgroups represented in the analysis with a single reference genome. The target branch of MKT is highlighted in each trio.

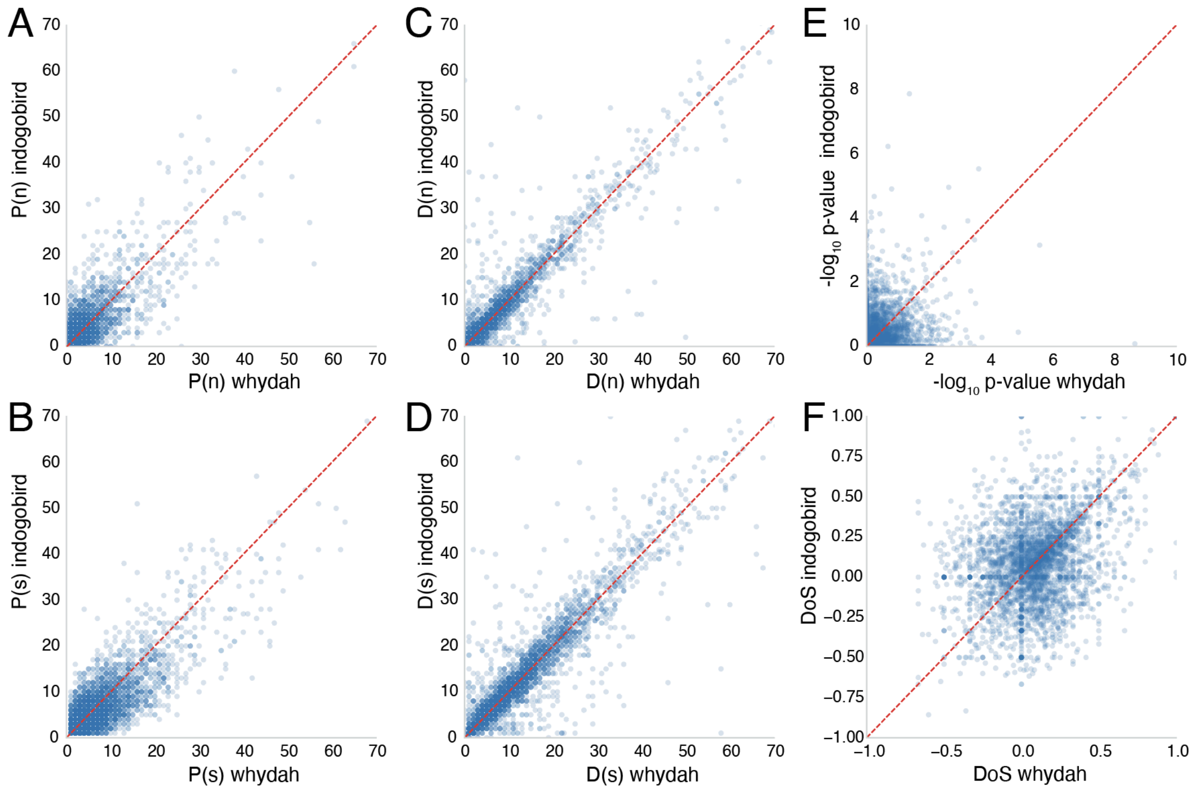


**Fig. S6. McDonald-Kreitman test (MKT) statistics compared between whydah and indigobird.**

(A-F): Each dot represents a gene in the coordinates of certain MKT statistics; (A) P(n): non-synonymous polymorphisms; (B) P(s): synonymous polymorphisms; (C) D(n): non-synonymous fixed differences; (D) D(s): non-synonymous fixed differences; (E) log10 p-value; (F) DoS: Direction of Selection.

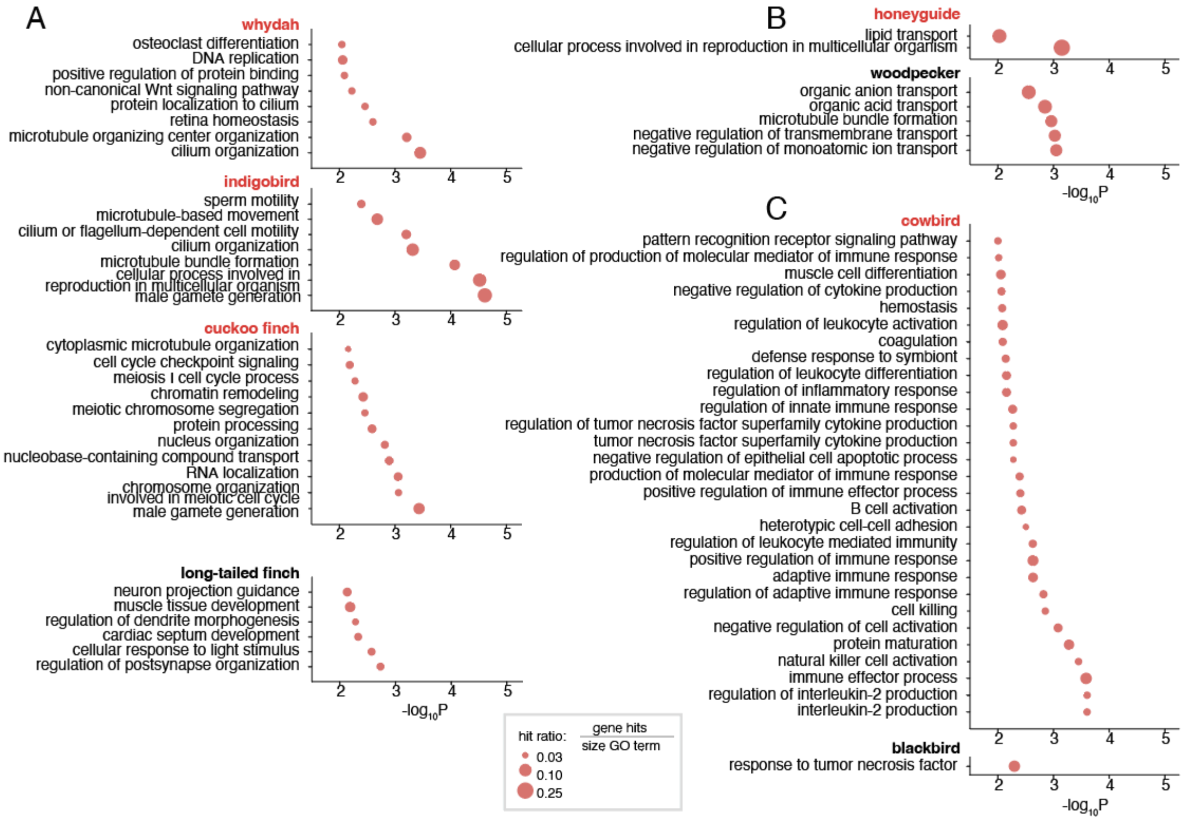


**Fig. S7. GO enrichment analysis of genes under positive selection in McDonald-Kreitman test.**
(A) Parasitic and estrildid finches clade: pin-tailed whydah, Cameroon indigobird, and cuckoo finch enrichments along with the outgroup long-tailed finch.
(B) Greater honeyguide with the outgroup downy woodpecker.
(C) Brown-headed cowbird with the outgroup red-winged blackbird.

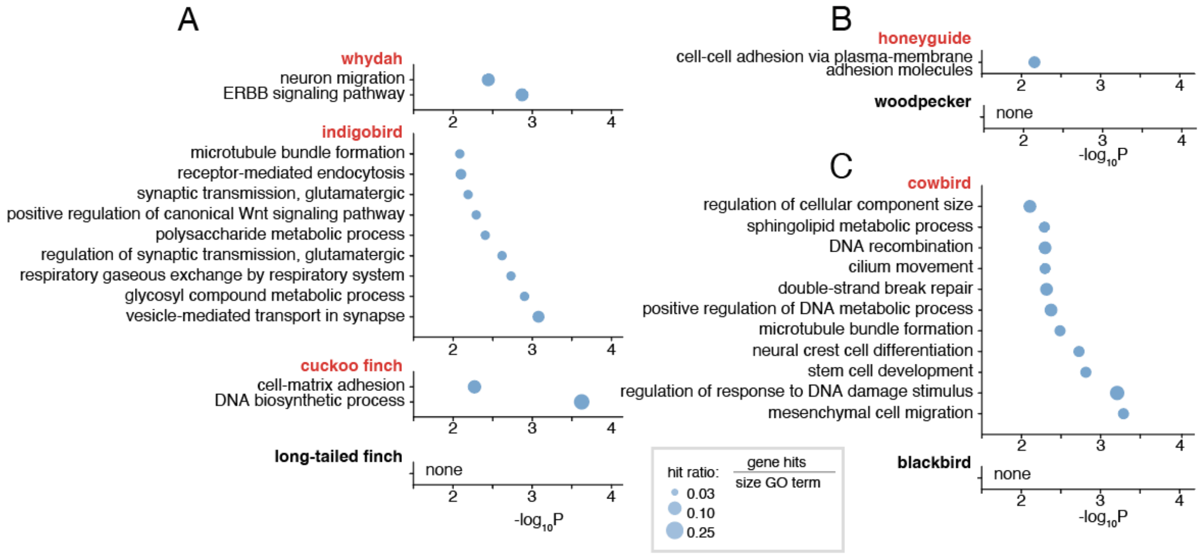


**Fig. S8. GO enrichment analysis of genes with negative direction of selection in McDonald-Kreitman test.**
(A) Parasitic and estrildid finches clade: pin-tailed whydah, Cameroon indigobird, and cuckoo finch enrichments along with the outgroup long-tailed finch.

(B) Greater honeyguide with the outgroup downy woodpecker.
(C) Brown-headed cowbird with the outgroup red-winged blackbird.

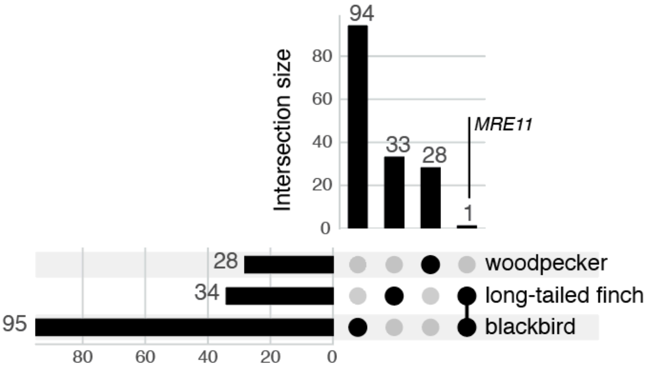


**Fig. S9. Genes under positive selection in outgroups in McDonald-Kreitman test.**
Upset plot illustrating total number of genes under positive selection (in MKT) in parental outgroups (horizontal bars on the left) along with the numbers of unique and overlapping genes (vertical bars).

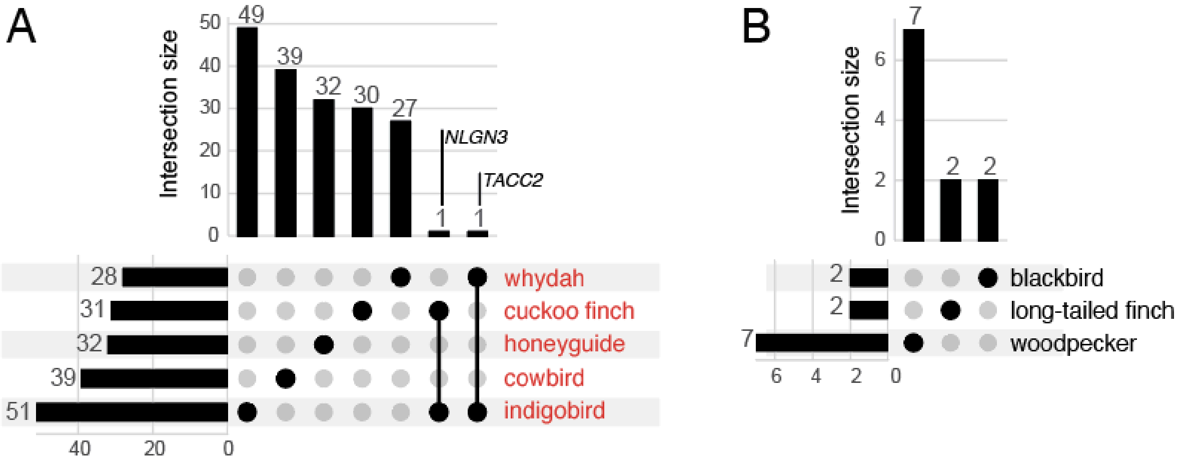


**Fig. S10. Genes with negative direction of selection in McDonald-Kreitman test.**
Upset plots showing the intersection of gene sets with negative direction of selection (DoS<0) in brood parasites (A) and parental outgroups (B).

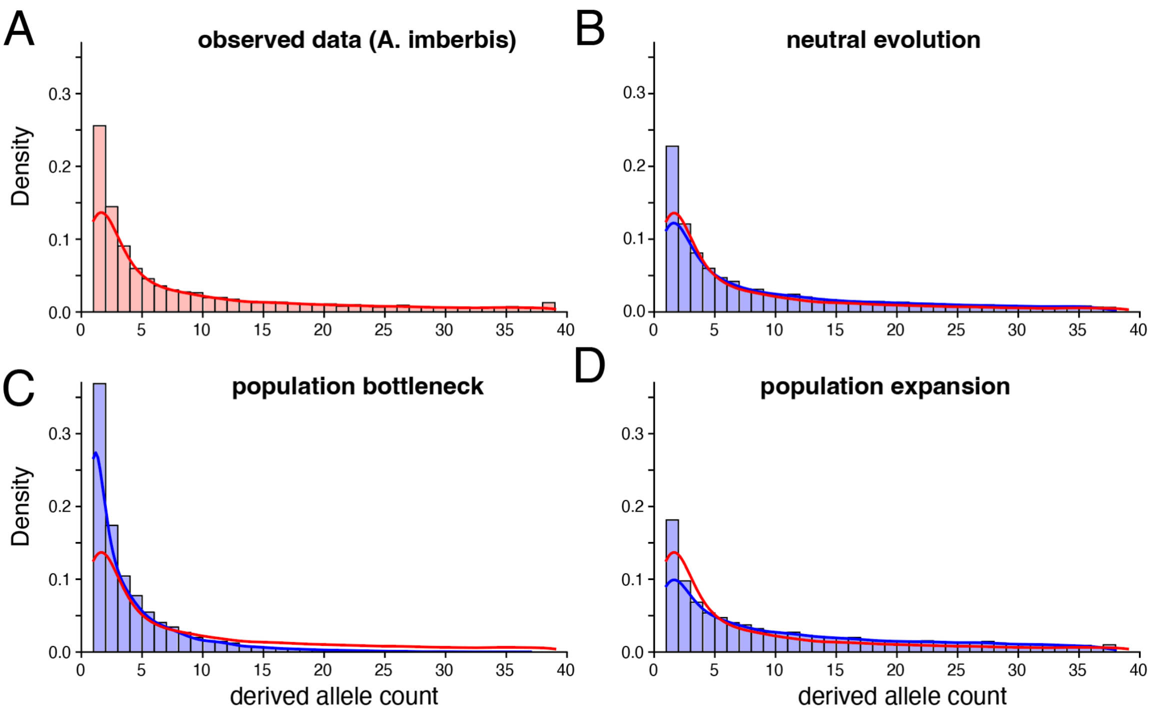


**Fig. S11. Site frequency spectra for MSPrime simulations compared to the cuckoo finch data.**
(A) Site frequency spectrum (SFS) of a 2-megabase (MB) fragment of the cuckoo finch scaffold CM062685.
(B-D) SFS of a 2MB genomic fragment simulated with MSPrime under different evolutionary scenarios: (B) neutral evolution; (C) recent population bottleneck (decrease in effective population size from 100,000 to 1000); and (D) recent population expansion (increase in effective population size from 100,000 to 200,000). The red line in simulation plots (B-D) repeats the density plots of the observed data from (A).

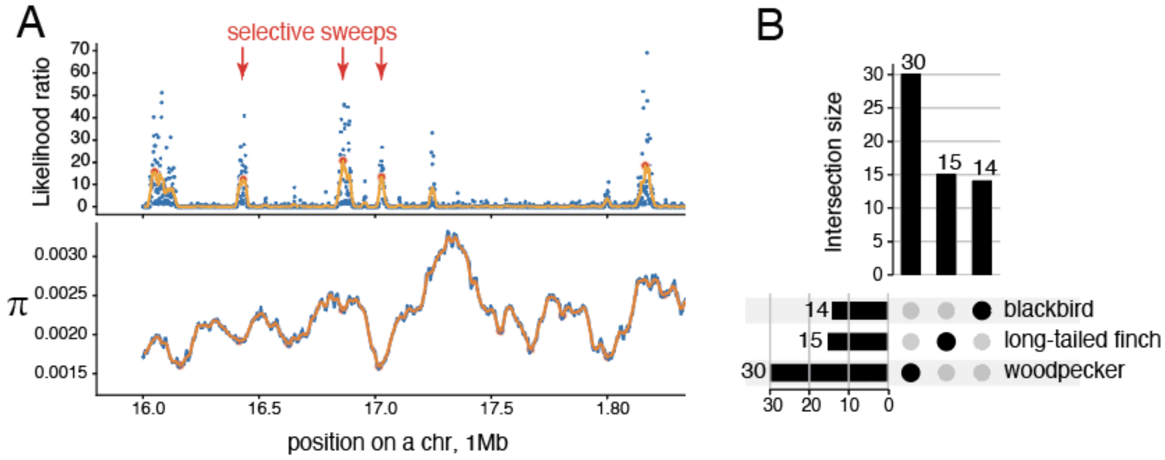


**Fig. S12. Selective sweeps in brood parasites and outgroups.**
(A) We identified genomic regions experiencing selective sweeps based on peaks in likelihood ratio from SweepFinder2 that overlap local minima in genetic diversity ($\pi$).
(B) Upset plot illustrating unique (horizontal bars on the left) and overlapping genes with associated selective sweeps in each parental outgroup species.
